# Post-traumatic osteoarthritis development is not modified by postnatal chondrocyte deletion of CCN2

**DOI:** 10.1101/2020.02.28.969105

**Authors:** Craig M Keenan, Lorenzo Ramos-Mucci, Ioannis Kanakis, Peter I Milner, Andrew Leask, David Abraham, George Bou-Gharios, Blandine Poulet

## Abstract

CCN2 is a matricellular protein involved in several critical biological processes. In particular, CCN2 is involved in cartilage development and in osteoarthritis. CCN2 null mice exhibit a range of skeletal dysmorphisms, highlighting its importance in regulating matrix formation during development, however its role in adult cartilage remains unclear. The aim of this study was to determine the role of CCN2 in postnatal chondrocytes in models of post-traumatic osteoarthritis (PTOA). CCN2 deletion was induced in articular chondrocytes of male transgenic mice at 8 weeks of age. PTOA was induced in knees either surgically or non-invasively by repetitive mechanical loading at 10 weeks of age. Knee joints were harvested, scanned with micro-CT, and processed for histology. Sections were stained with toluidine blue and scored using the OARSI grading system. In the non-invasive model cartilage lesions were present in the lateral femur but no significant differences were observed between wildtype (WT) and CCN2 knockout (KO) mice 6 weeks post-loading. In the surgical model, severe cartilage degeneration was observed in the medial compartments but no significant differences were observed between WT and CCN2 KO mice at 2, 4, and 8 weeks post-surgery. We conclude that CCN2 deletion in chondrocytes did not modify the development of PTOA in mice, suggesting that chondrocyte expression of CCN2 in adults is not a critical player in protecting cartilage from the degeneration associated with PTOA.

**Summary Statement:** Post-natal deletion of CCN2 in chondrocytes does not affect the development of post-traumatic osteoarthritis in mice.

## Introduction

Osteoarthritis (OA) is a major chronic degenerative disease of the joint, with limited therapies available to inhibit or slow disease progression. Mechanical trauma represents a major risk factor for OA with mouse models of post-traumatic OA (PTOA) widely used to study OA development. These include surgical models for various OA severity development (Kamekura et al., 2005) and non-invasive repetitive joint trauma model including the one established by Poulet *et al* (Poulet et al., 2011). Both models show similar hallmarks as seen in human OA, including progressive articular cartilage degradation, subchondral bone sclerosis, osteophyte formation and synovial fibrosis and activation. In this study, we used both a surgical model and a non-invasive model to determine the role of the matricellular protein CCN2 in severe and moderate PTOA severities.

CCN2, also known as Connective tissue growth factor (CTGF), is a matricellular protein involved in key cellular functions including proliferation, adhesion and differentiation (Kubota and Takigawa, 2015), along with several complex biological processes including chondrogenesis (Perbal, 2004). Deletion of CCN2 during development demonstrated its essential role as a regulator of skeletal development by promoting endochondral ossification through the proliferation and differentiation of growth plate chondrocytes (Shimo et al., 2000, Takigawa et al., 2003). In addition, CCN2 has been suggested as a potential cartilage repair factor (Nishida et al., 2004); continuous cartilage-specific overexpression of CCN2 has been shown to protect joints from age-related OA development, highlighting a chondro-protective role of CCN2 (Itoh et al., 2013).

CCN2 is expressed in adult articular cartilage, albeit at low levels (Tang et al., 2018, Nishida et al., 2004), and its expression significantly increases in OA chondrocytes (Omoto et al., 2004). However, the role of chondrocyte-specific expression of CCN2 in OA is still unknown. There is evidence that CCN2 may be involved in the pathogenesis of OA as CCN2 overexpression in the synovial lining of mouse knee joints resulted in the development of transient fibrosis and cartilage damage (Davidson et al., 2006). In an inducible KO model, global deletion of CCN2 postnatally resulted in protection from surgically induced OA (Tang et al., 2018), which suggests a negative role for CCN2 in OA. These contradictory roles of CCN2 in joint health and OA clearly need to be further explored. Therefore, to understand the importance of CCN2 in adult cartilage, we examined the effect of postnatal CCN2 deletion specifically in chondrocytes using two models of PTOA.

## Materials and Methods

### Animals

All work was carried out in accordance with the UK Home Office guidelines and regulations under the Animals (Scientific Procedures) Act 1986. All mice (C57CBA background) were housed in the specific pathogen free biological services unit at the University of Liverpool, UK and housed in cages of up to 5 mice, with 12h light/dark cycle, and ad libitum food and drink. Using chondrocyte specific aggrecan (Acan) enhancers, two conditional CCN2 KO mouse models were generated. The first contained an Acan -10kb CreER^T2^ enhancer as described by Han and Lefebvre (Han and Lefebvre, 2008). This produced Acan -10kb CreER^T2^ x CCN2^fl/fl^ mice in which CCN2 was deleted from articular chondrocytes. The second contained an Acan -30kb CreER^T2^ enhancer described by Li *et al* (Li et al., 2018). This produced Acan -30kb CreER^T2^ x CCN2^fl/fl^ mice where CCN2 was deleted from all chondrocytes. Deletion of CCN2 was regulated using a tamoxifen inducible CreER^T2^.

### Tamoxifen induction of Acan CreER^T2^

Prior to the start of all *in vivo* experiments, deletion of CCN2 or STOP in reporter td Tomato using Acan CreER^T2^ was induced at 8 weeks of age using tamoxifen (Sigma Aldrich, UK). All mice, whether WT or CCN2^fl/fl^, were administered tamoxifen intraperitoneally at a dose of 1mg/10g body weight on days 1, 3, and 5, and were weighed prior to injection on each day. Following tamoxifen injections, mice were left for 1 week before any experimental work commenced. Td tomato mice were sacrificed four weeks after last tamoxifen injection.

### Non-invasive mechanical loading model of PTOA

Right knees of 10 week-old male Acan -30kb CreER^T2^ x CCN2^fl/fl^ (n = 13 (Cre^WT^), n = 14 (Cre^+/o^)) mice were loaded non-invasively, using a model previously describe*d* (Poulet et al., 2011) to induce PTOA. Briefly, mice were anaesthetised and the right leg placed in custom-made cups with the knee in flexion. A peak load of 9N was applied for 0.05 seconds, with a rise and fall time of 0.025 seconds, and a baseline hold time of 9.9 seconds for 40 cycles. A baseline load of 2N was employed to keep the tibia in place during peak loading. All mice were subjected to this pattern 3 times per week for 2 weeks. Loading was performed using an ElectroForce 3100 (TA Instruments, USA). Mice were weighed after each loading episode, and on a weekly basis following completion of the loading regimen. All mice were sacrificed 6 weeks post-loading and samples prepared for micro-CT and histological analysis.

### Surgical model of PTOA

Left knee joints of 10-week-old Acan -10kb CreER^T2^ x CCN2^fl/fl^ mice underwent surgical transection of the medial meniscus (MM) and the medial meniscotibial ligament (MMTL). Mice were induced and maintained under a plane of general anaesthesia using isoflurane during the surgical procedure.. A small incision was made over the medial aspect of the patella tendon and the joint capsule incised. Using blunt dissection small amounts of fat were removed allowing for visualisation of the MM and MMTL. Using a scalpel, the MM and MMTL were transected using an upwards motion from the cranial horn of the MM on the proximal tibial plateau. Once transected the joint capsule and the skin were sutured. Mice were immediately transferred to a heated post-operative recovery room. They were monitored daily to ensure they were in good health. Mice were sacrificed 2 weeks post-op (n = 4 (Cre^WT^), n = 14 (Cre^+/o^)), 4 weeks post-op (n = 7 (Cre^WT^), n = 7 (Cre^+/o^)), and 8 weeks post-op (n = 7 (Cre^WT^), n = 8 (Cre^+/o^)). Samples were prepared for micro-CT and histological analysis.

### Specimen preparation

All animals were sacrificed by cervical dislocation. Experimental and contralateral joints were dissected, immediately fixed in 10% neutral buffered formalin for 24hrs and transferred to 70% ethanol for storage.

### Micro-CT analysis

Experimental and contralateral joints were scanned at a resolution of 4.5µm using a 0.25mm aluminium filter, with a rotation step of 0.6° (Skyscan 1272, Bruker microCT, Belgium). Image reconstruction was performed using NRecon software (Bruker microCT, Belgium), followed by manual selection of regions of interest for tibial and femoral epiphysis, and joint space (including menisci). Bone volume/tissue volume (BV/TV) and tissue volume (TV) were determined. Data were tested for normality and Student’s t-test was used for statistical evaluation with significance set at P<0.05.

### Histological analysis

Samples were decalcified in either 10% ethylenediaminetetraacetic acid (EDTA) (Sigma Aldrich, UK) for 2 weeks or 10% formic acid (Sigma Aldrich, UK) for 1 week. Once decalcified, samples were given a processing number independent of their genotype, processed, paraffin embedded in either the coronal (surgical model) or sagittal plane (loading model), and 6µm sections were taken throughout the entire joint. Sections across the joint at 120µm intervals were stained with toluidine blue/fast green (0.04% in 0.1M sodium acetate buffer, pH 4.0) and cartilage lesion severity graded using the OARSI histopathology initiative scoring method (Glasson et al., 2010). Grading each of the four compartments of the tibio-femoral joint (lateral and medial tibia and femur) throughout the entire joint allowed for the determination of a maximum lesion grade (most severe lesion) for the whole joint and each individual compartment. The mean score, which involved determining the average grade across multiple slides was calculated for each joint and each compartment (Poulet et al., 2011). The summed score was determined by adding together the maximum score of each compartment per joint. Osteophyte formation was graded histologically using the scale described by Kamekura *et al* (Kamekura et al., 2005). Statistical analysis was performed using a Mann-Whitney test. Data are presented as box plots of interquartile range, median, minimum and maximum and showing all individuals.

### Cryosectioning

Samples were dissected, fixed in 4% paraformaldehyde at 4 °C for 24hrs, decalcified in 10% EDTA for 2 weeks, embedded using OCT embedding media (Tissue-Tek, Sakura Europe) in either the coronal (WT) or sagittal (Cre^+/o^) plane, and stored at -80°C until required. Sections were taken at 5µm until the middle of the joint was reached. A section showing the entire tibio-femoral joint was then collected and stained with Hoechst stain for 30 mins. Images were obtained using a Zeiss Axio Observer apotome microscope (Zeiss, Germany).

## Results

### CCN2 was successfully deleted in chondrocytes in adult mice

To verify that deletion of CCN2 in chondrocytes had occurred in response to tamoxifen injection, a tail-tip was taken post-cull from each individual mouse, and PCR performed on extracted genomic DNA for genetic recombination and exon deletion. All mice contained a 1000bp band corresponding to the floxed-CCN2 allele, and Cre^+/o^ mice contained an additional band at around 500bp corresponding to the allele generated by Cre-recombination resulting in the Cre-mediated deletion of CCN2 (Fig. 1). Cre^WT^ mice showed no recombination. To validate the efficiency of the CreER^T2^ system and the subsequent deletion of the transgene in these new Acan -30kb CreER^T2^ mice, CCN2^fl/fl^ with Cre^+/o^ (CCN2 KO) mice were crossed with a tdTomato reporter mouse, which enabled detection of recombined CreER^T2^ via fluorescence, following administration of tamoxifen to the mice. In control corn oil treated mice, no tdTomato was seen in the whole joint cells (only a chondrocyte was positive; Fig. 1), demonstrating no recombination of CreER^T2^ occurred. Following tamoxifen treatment, there was clear recombination of CreER^T2^ as evident by the intense expression of tdTomato protein in all chondrocytes located in both the articular cartilage and the growth plate (Fig. 1).

**Figure 1:**
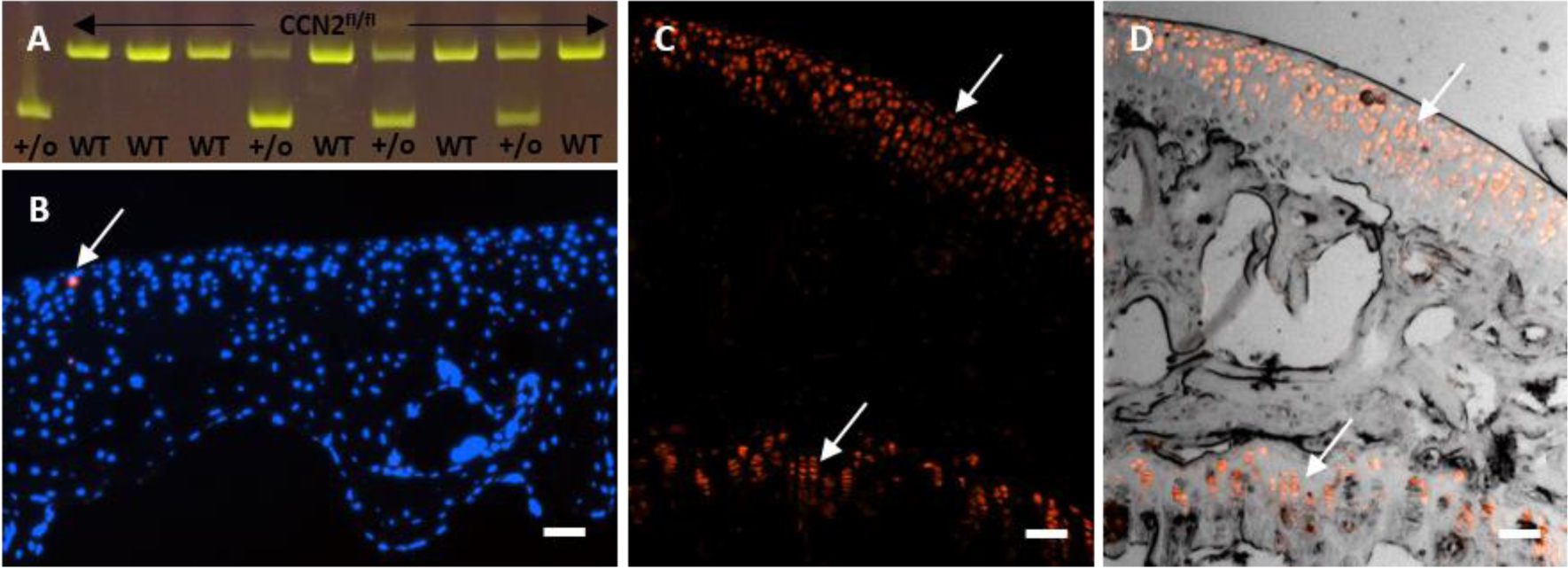
Confirmation of *Cre* recombination and CCN2^fl/fl^ transgene deletion in chondrocytes in -30kb *Acan* CreER^T2^ adult mice. (A) Genotyping from tail tips genomic DNA from WT and CCN2 Cre^+/o^ mice showed the presence of the CCN2 flox product in all mice (first band) and the additional lower band in Cre^+/o^ mice only (+/o) around 500bp in size confirming Cre recombination and deletion of the CCN2 floxed product. (B, C, & D) Histological images of the tibial epiphysis in CCN2 Cre^+/o^ crossed with the tdTomato reporter mouse 4 weeks after the last corn oil control (B) or tamoxifen injection (C-D). Corn oil treatment showed no recombination (except one single chondrocyte red; arrow B; blue staining of nuclei with Hoechst stain). (C-D) Tamoxifen treatment showed expression of tdTomato fluorescence (red) in chondrocytes throughout the articular cartilage and growth plate (top and bottom arrows respectively). Scale bar = 100µm.

### Conditional CCN2 deletion in chondrocytes does not prevent the development of osteoarthritis in a non-invasive loading model of trauma-induced OA

Mice (CCN2 KO and WT) were treated with tamoxifen then mechanically loaded to induce PTOA. Histological examination of the loaded limb in both WT (n=13) and CCN2 KO (n=14) mice showed the development of OA lesions with loss of hyaline articular cartilage and exposure of the articular calcified cartilage primarily in the lateral femur (Fig. 2). Assessment of the severity of the lesions in each compartment throughout the entire tibio-femoral joint (medial and lateral, tibia and femur) showed no significant differences in the AC lesion mean, maximum, and summed severity scores between WT and CCN2 KO mice. MicroCT analysis of the lateral femur and tibia epiphyseal bone showed no differences in BV/TV between WT and KO (Fig. 2). Together, these data indicate that deletion of CCN2 postnatally from aggrecan-expressing chondrocytes had no effect on the development of OA in a non-invasive model of moderate PTOA.

**Figure 2:**
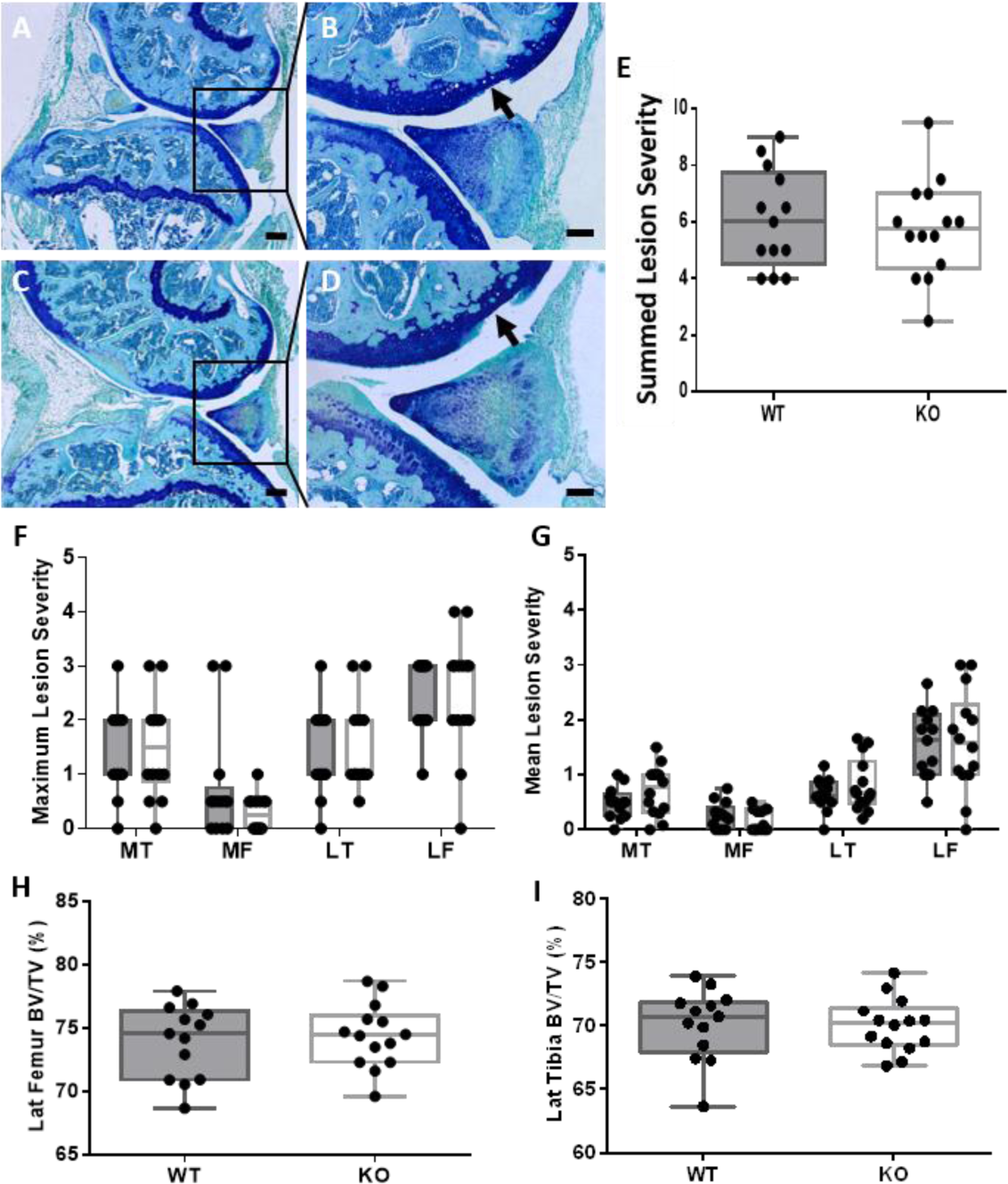
Deletion of CCN2 in chondrocytes does not prevent OA development in a non-invasive loading model of PTOA. (A-D) Toluidine blue stained sections of the tibio-femoral joint of WT (A,B) and CCN2^fl/fl^ KO (C,D) mice showed development of AC lesions localized to the lateral femur in loaded knee joints (arrowed). (E-G) AC lesion severity scores across the whole knee joint showed no differences between WT and CCN2^fl/fl^ KO mice in (E) summed maximum scores nor in (F) maximum and (G) mean lesion severity scores for each joint compartment. (H, I) BV/TV of the epiphyseal bone in the lateral femur (H) and lateral tibia (I) showed no significant difference in bone structure between WT and CCN2^fl/fl^ KO. (MT = medial tibia, MF = medial femur; LT = lateral tibia; LF = lateral femur). Scale bar = 200µm (A,C) and 100µm (B,D).

### CCN2 deletion specifically in chondrocytes does not prevent the development of osteoarthritis in a surgical model of severe PTOA

To confirm that deletion of CCN2 in chondrocytes of adult mice had no effect on OA, a second model of PTOA was used. This model incorporated a different Cre (Acan -10kb CreER^T2^) which had been previously shown to express in articular chondrocytes (Han and Lefebvre, 2008). At 2 weeks post-surgery, moderate AC lesions were already visible in the medial compartment of both WT and CCN2 KO mice with exposure of the underlying calcified cartilage (Fig. 3). At 4 weeks post-surgery, the damage was extensive with both WT and CCN2 KO suffering substantial loss of articular cartilage in the medial compartment of the tibio-femoral joint. By 8 weeks post-surgery there was widespread, significant damage across the entire medial side of the tibio-femoral joint in both WT and CCN2 KO mice, including near-complete loss of AC. Cartilage lesion scoring showed no significant differences between WT and CCN2 KO at all time-points.

**Figure 3:**
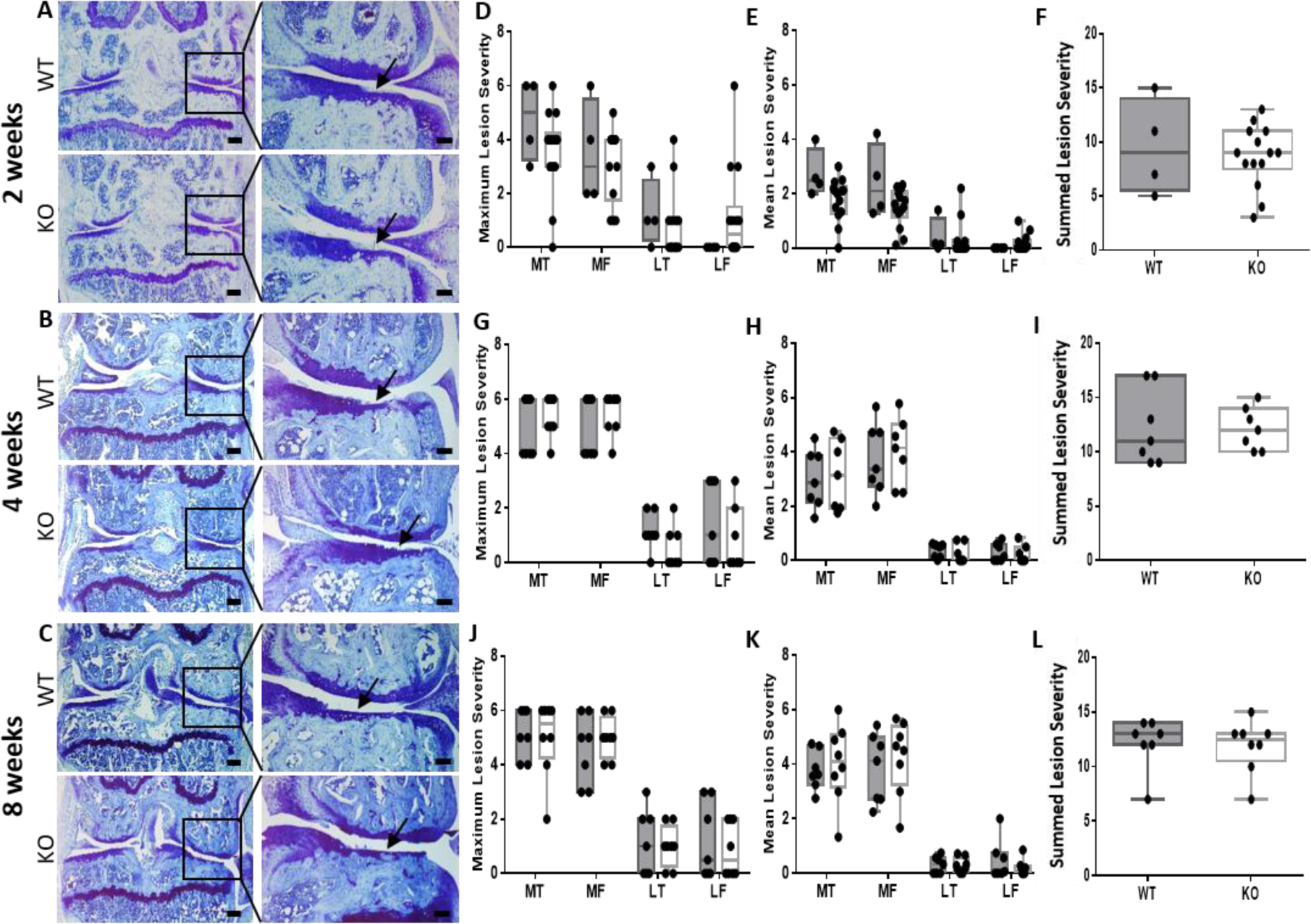
Deletion of CCN2 in chondrocytes does not prevent AC lesion severity in a surgical model of severe PTOA. (A-C) Toluidine blue stained sections from WT and CCN2^fl/fl^ KO mice 2, 4 and 8 weeks post-surgery showed development of OA on the medial tibia (arrows) in both WT (top panel) and CCN2^fl/fl^ KO (bottom). (D-L) Maximum and mean lesion severity in each individual joint compartment and summed maximum scores showed no significant difference between OA severity in WT and CCN2^fl/fl^ KO mice at (D-F) 2 weeks, (G-I) 4 weeks, and (J-L) 8 weeks post-surgery. (MT = medial tibia, MF = medial femur; LT = lateral tibia; LF = lateral femur). Scale bar = 200µm (A, B, & C left panels), 100µm (A, B, & C right panels).

Osteophyte formation was observed at 2 weeks post-surgery in WT and CCN2 KO mice (Fig. 4) and advanced to fully ossified osteophytes at 4- and 8-weeks post-surgery (Fig. 4), with no significant differences observed between WT and CCN2 KO mice at all time-points. Data generated from the surgical injury model of OA suggest that deletion of CCN2 in chondrocytes postnatally shows no difference in PTOA development

**Figure 4:**
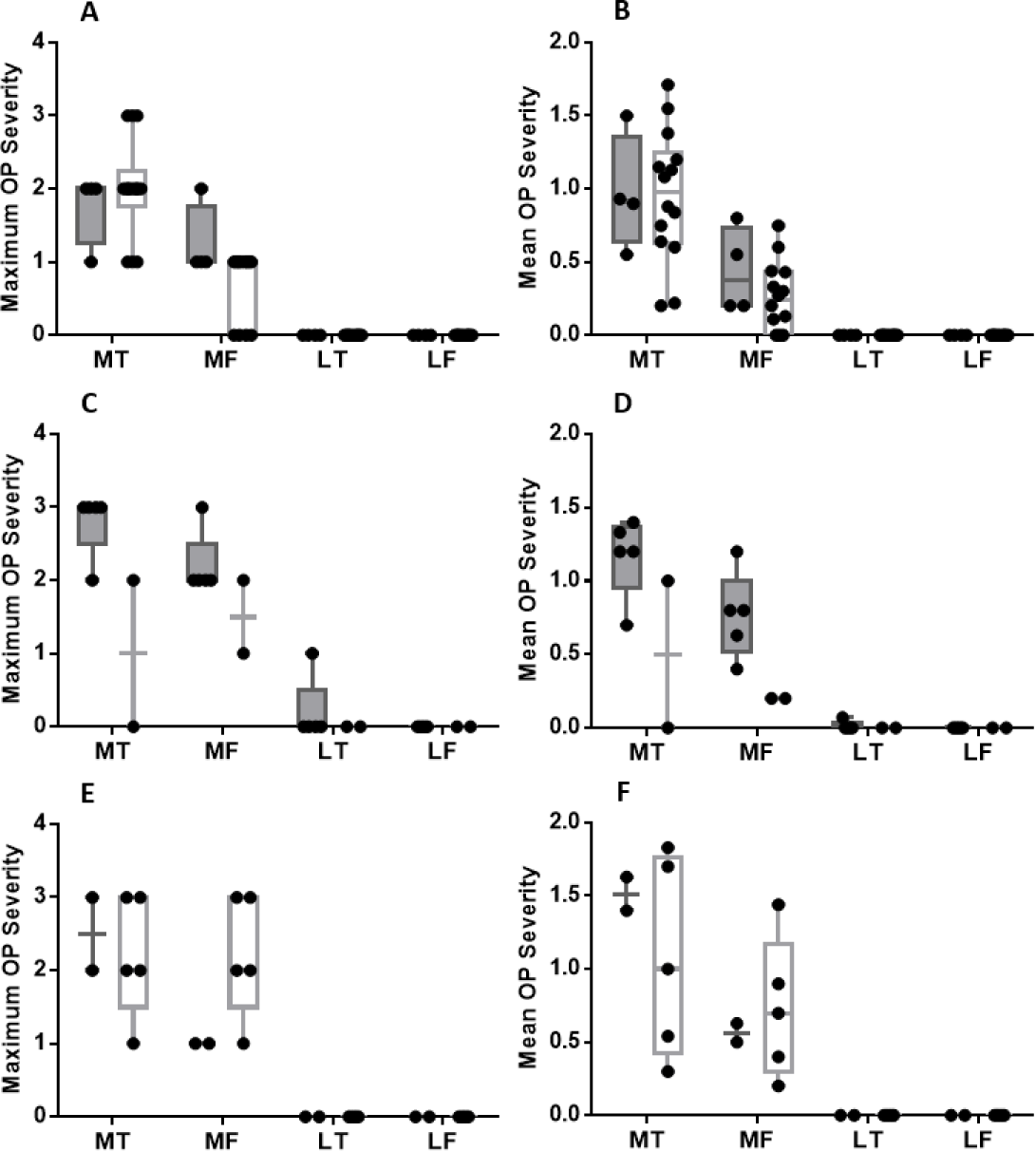
Deletion of CCN2 in chondrocytes does not affect osteophyte maturity in a surgical model of PTOA. Maximum and mean osteophyte severity scores across all knee joint compartments showed no significant differences between WT and CCN2^flf/fl^ KO mice at 2 weeks (A-B), 4 weeks (C-D) and 8 weeks (E-F) post-surgery.

## Discussion

This study aimed to determine the postnatal role of CCN2 in articular cartilage during PTOA in two different models. We have found that despite its importance during skeletal development and its increased expression by OA chondrocytes, CCN2 in articular cartilage chondrocytes in adult joints did not play a significant role in preventing PTOA development.

A number of studies have focused on the regenerative ability of CCN2 due to its role in cellular proliferation and differentiation (Takigawa et al., 2003, Shimo et al., 2000). Its role in chondrogenesis highlighted its potential as a possible regenerative therapy for the treatment of OA, particularly as exogenously added rCCN2 did not stimulate hypertrophy of chondrocytes *in vitro* (Nishida et al., 2002). Moreover, treatment of articular cartilage defects with rCCN2 showed repaired cartilage to be structurally similar to healthy articular cartilage (Nishida et al., 2004). A recent study by Tang and colleagues showed that a postnatally induced global deletion of CCN2 caused a thickening of cartilage that resulted in protection from ageing-induced OA (Tang et al., 2018). Since postnatal global deletion of CCN2 impeded OA development, it was hypothesised that postnatal chondrocyte-specific deletion of CCN2 would also protect cartilage from the development of OA following trauma.

The lack of effect of chondrocyte-specific compared to global deletion of CCN2 in mice (which protected from OA development) suggests that CCN2 expression in other tissues, including those surrounding the tibio-femoral joint, may be required for protection from OA. Indeed, CCN2 is expressed by a number of other cells/tissues in the joint such as synovial fibroblasts (Davidson et al., 2006) and has been shown to induce OA hallmarks such as synovial fibrosis, cartilage degeneration and osteophyte formation. It is also entirely feasible that the lack of effects seen in chondrocyte-specific deletion in our studies is linked to secretion of CCN2 from these other joint tissues and its release into the joint space, thereby preventing any effects of chondrocyte-specific deletion on OA development from being observed. Other cells, including those derived from mesenchymal stem cells (MSCs) and MSC-like progenitor cells, may have also affected the levels of CCN2 in the joint. These cells are present in all joint tissues and can differentiate into several populations of cells including those with chondrogenic potential (Barry and Murphy, 2013). These cells, which are not targeted for CCN2 deletion, may have been recruited in response to lesion formation, leading to increased expression of CCN2 in the joint and subsequent loss of the KO effect on OA development.

It should be noted that the absence of any observable responses in our studies is not as a result of insufficient Cre recombinase activity and the subsequent deletion of CCN2 from chondrocytes. The efficiency of Cre recombinase activity for the -10kb Acan enhancer had been tested previously and shown to be effective (Cascio et al., 2014), while the new -30kb enhancer showed equally greater number of chondrocytes expressing the transgene (Li et al., 2018). To confirm the efficiency of this new Acan specific CreER^T2^ system, Acan -30kb CreER^T2^ with CCN2 ^fl/fl^ mice were crossed with tdTomato reporter mouse, which allowed for detection of Cre recombinase activity following tamoxifen injection by visualisation of the tomato fluorescence compared to oil injected control, and confirmed recombination and hence CCN2 deletion, from all articular cartilage chondrocytes. Furthermore, the use of two different Acan-Cre systems to drive Cre-recombinase expression in chondrocytes ensured any responses were independent of Cre activity targeting and were most likely a direct result of the action of CCN2 deletion.

In this study, we used two models of PTOA, with different severities of OA progression. The non-invasive loading of the knee has previously been shown to induce moderate OA lesions on the lateral femur (Poulet et al., 2011), whereas surgical intervention is known to lead to a higher degree of OA severity. The importance of using different models pertain to the fact that, although similar pathologies can be seen in both models, both may trigger different cellular responses linked to the severity of the disease. For example, the surgical model might trigger more severe inflammatory responses. In addition, the mechanical environment is severely affected in the surgical model throughout the whole study, whereas the traumatic loads are applied at specific and controlled times, with a maximum of 6 episodes of 7 minutes; the rest of the time, the mechanical environment is relatively normal compared to that engendered by surgical instability. In future studies, the lack of effects of chondrocyte specific CCN2 expression in adults could be tested in other models of OA, including ageing and obesity-induced OA.

In conclusion, this study showed through the use of two models of trauma-induced OA that CCN2 expression by chondrocytes is not required for maintenance of cartilage in adults and that CCN2 expression by other tissues within the joint may be more important for any effect to be observed.

## Author contributions

1. Study conception and design: BP, GBG, DA; Acquisition of data: CK, LRM, IK, PM, AL, GBG, BP. Data analysis and interpretation: CK, LRM, IK, PM, AL, DA, GBG, BP; Obtaining of funding: BP, GBG, DA.
2. Drafting and revision of manuscript: all authors have contributed to the draft and revision of the manuscript and their comments have been added to the final version when appropriate.
3. Final Approval of the Manuscript: all authors have reviewed the final version of the manuscript and approved the version to be published.

